# *In vivo* quantification of creatine kinase kinetics in mouse brain using ^31^P-MRS at 7 Tesla

**DOI:** 10.1101/2024.09.09.611986

**Authors:** Mohamed Tachrount, Sean Smart, Jason Lerch, Antoine Cherix

## Abstract

^31^P-MRS is a method of choice for studying neuroenergetics *in vivo*, but its application in the mouse brain have been limited, often restricted to ultra-high field (>7 Tesla) MRI scanners. Establishing its feasibility on more readily available preclinical 7 Tesla (T) scanners would create new opportunities to study metabolism and physiology in murine models of brain disorders. Here, we demonstrate that the apparent forward rate constant (k_f_) of creatine kinase (CK) can be accurately quantified using a progressive saturation-transfer approach in the mouse brain at 7T. We also find that a reduction of approximately 20% in the breathing rate of anesthetized mice can lead to a 36% increase in k_f_ attributable to a drop in intracellular pH and mitochondrial ATP production. To achieve this, we used a test-retest analysis to assess the reliability and repeatability of ^31^P-MRS acquisition, analysis and experimental design protocols. We report that most ^31^P-containing metabolites can be reliably measured using a localized 3D-ISIS sequence, which showed highest SNR amplitude, SNR consistency and minimal T_2_ relaxation signal loss. Using this protocol, our study identifies, for the first time, key physiological factors influencing mouse brain energy homeostasis *in vivo* and provides a methodological basis that will guide future studies interested in implementing ^31^P-MRS on preclinical 7T scanners.

## Introduction

Phosphorous Magnetic Resonance Spectroscopy (^31^P-MRS) is a well-established and time-tested method for measuring energy metabolism in vivo. This technique allows for the quantification of phosphate-containing molecules involved in energy homeostasis, such as phosphocreatine (PCr), adenosine triphosphate (ATP), inorganic phosphate (Pi) and nicotinamide adenine dinucleotide (NADH and NAD^+^). Additionally, other phosphate-containing molecules can be observed, including precursors and intermediates of phospholipid metabolism^1^, such as phosphoethanolamine (PE) and phosphocholine (PC), both often referred to as phosphomonoesters (PME), or glycerophosphoethanolamine (GPE) and glycerophosphocholine (GPC), known as phosphodiesters (PDE). Moreover, intracellular pH can be indirectly measured from the chemical shift differences of Pi and PCr^2^. Finally, when combined with a saturation transfer preparation, ^31^P-MRS provides a unique opportunity to measure the activity rates of key enzymes like creatine kinase (CK) and ATP synthase^3^, both of which are critical in regulating cellular ATP production.

The foundations of ^31^P-MRS were established over four decades ago, with the method tested and validated across various samples and organisms, including yeast^4^, perfused rodent heart^5,6^, rodent brain^7–9^ and, ultimately, human heart^10^ and brain^11,12^. Since then, ^31^P-MRS has been refined and applied in numerous studies to investigate the metabolic basis of various disorders in humans^13,14^ and in animal models^15^.

Despite the high potential of ^31^P-MRS for studying *in vivo* metabolism, research on brain energetics in animal models has been scarce and often restricted to ultra-high field (> 7 Tesla) scanners. Acquiring ^31^P-MRS spectra from small volumes like the rodent brain is technically challenging, making ultra-high magnetic field scanners a valuable tool for enhancing signal-to-noise ratio (SNR) and spectral resolution. Although some studies using lower field scanners (< 7T) have been attempted in mice, reporting metabolite changes in the brains of tumor-bearing animals^16–18^, in a scrapie model^19^ or following overexpression of astroglial growth hormone^20^, these studies have often been limited by their localization protocols and have primarily quantified only the metabolite peaks with the highest signals, such as PCr, ATP or occasionally Pi, with limited or no information on the reliability of the measured signals.

Ultra-high field scanners (e.g. 9.4T or 14T) have improved sensitivity and spectral resolution, enabling the study of neuroenergetic alterations in mouse models of neurodegenerative^21–24^, neurological^25,26^ or neuropsychiatric^27^ disorders. This higher resolution has made it possible to acquire data on more complex biochemical processes, such as CK rate^22^ measurements, or to distinguish between important metabolic pools, such as intracellular versus extracellular Pi or NADH versus NAD+^27^. While the advantages of higher field scanners for ^31^P-MRS are well-established, including shorter T1 times and improved chemical shift separation^28^, their prohibitive costs and technical challenges - such as B0 and RF inhomogeneities, especially with surface coils - have restricted their use to niche applications. Consequently, most recent method developments in mouse brain imaging have solely focused on using 9.4T scanners^29–31^.

Preclinical 7T scanners offer a balanced compromise between cost-effectiveness and imaging performance, providing sufficient resolution and sensitivity for many applications while remaining more accessible and less prone to high-field-related artifacts. 7T scanners require lower initial investment and operational costs, need less specialized infrastructure, and present fewer technical challenges due to their lower field strength, making them more widely available. However, the feasibility and repeatability of ^31^P-MRS measurements in the mouse brain using such scanners are less well understood. In this study, we explore the potential of using ^31^P-MRS at 7T to measure brain energy metabolism in the mouse brain, evaluating its benefits and reliability. We compare three standard sequences, assess the reliability and repeatability of the resulting metabolite quantifications, and examine the feasibility of using saturation transfer to quantify creatine kinase rates *in vivo*.

## Materials and Methods

### Animals

Male and female adult C57/BL6J mice obtained from in-house breeding were housed in standard IVC cages in a normal 12h day-light cycle environment at a temperature of 20-24°C and humidity of 46-65%. Animals had *ad libitum* access to standard rodent chow diet and water. All experiments were carried out with approval by the UK Home Office, under the Animals (Scientific Procedures) Act 1986, and in compliance with the ARRIVE (Animal Research: Reporting *in vivo* Experiments) guidelines.

### Experimental design

#### Test-retest analysis of ^31^P-MRS

Six adult male C57BL/6J mice (35±3g, 33-weeks old) were scanned using ^31^P-MRS on two occasions at one week interval between scans. A test-retest analysis was done using three different sequences by comparing their respective signal-to-noise ratio (SNR) and the coefficients of variation (CVs) of their SNRs. The three sequences were applied in random order, but that order kept the same for each mouse on the two scan repetitions. Then, the performance of the best sequence was assessed by reporting the quantification error (Cramer-Rao lower bound; CRLB) of each metabolite component for increasing acquisition times. This increase in acquisition time, from t_0_ to t_f_, was estimated by gradually increasing the number of included averages in random order from an initial scan of duration t_f_. The number of included averages is thus reported in time units for clarity throughout the manuscript. The CVs of the most reliable metabolite quantifications (PCr/ATP and Pi/ATP) were then determined for between-group, between-session and within-session comparisons. The resulting CVs were ultimately used to perform power calculations for subsequent studies.

#### Group comparison for saturation-transfer (ST) ^31^P-MRS

Nine adult male (32±2g) and eleven female (23±2g) C57BL/6J mice (16-weeks old) were scanned using ST ^31^P-MRS. The estimated Creatine Kinase forward rate constant (k_f_) was then correlated to physiological, biological and acquisition-related parameters. Animals were then grouped based on the biological parameter with strongest correlation coefficient, using the median as a separation cutoff for the two groups.

### *In vivo* MRI acquisition

Animals’ anesthesia was induced with 2-3% isoflurane in air mixture containing 30% O_2_. Animals were monitored during the entire scan for physiological parameters using a small animal monitoring system (SA Instruments Inc., New York, USA). Breathing rate per minute was maintained between 75 – 105 rpm by adjusting the isoflurane dose between 1 and 1.5%. Rectal temperature was kept at 36.8±0.5°C with a circulating heating water bath and assessed using a temperature rectal probe. Animals were scanned using a 7T (70/20) BioSpec MRI scanner (Bruker, Ettlingen, DE) using Paravision 360.1.1. A dual tuned ^31^P/^1^H surface coil with a single 10mm ^31^P loop (PulseTeq Ltd, Chobham, UK) was used as transceiver (Tx/Rx). A set of anatomical T_2_-weighted images were acquired in axial, sagittal and coronal orientations (TurboRARE, TE/TR= 11ms/2632.6ms, RARE-factor=8) for localization and voxel placement for subsequent magnetic resonance spectroscopy (MRS) acquisitions.

### *In vivo* ^31^P- Magnetic Resonance Spectroscopy (^31^P-MRS)

For ^31^P-MRS, a 160uL voxel (8×5×4mm^3^) was placed in a brain area enveloping both hippocampus and hypothalamus. Shimming (MAPSHIM) was performed in the voxel to reach a water Full Width at Half Maximum (FWHM) of 26±2Hz. The acquisition was performed using Point-Resolved Spectroscopy (PRESS)^32^, semi-Localized by Adiabatic Selective Refocusing (sLASER)^33^ and Image-Selected In Vivo Spectroscopy (3D-ISIS)^34^ in randomized order with similar acquisition parameters (Npoints=1024, AcquisitionBW=40ppm, TR=4s, Averages/Repetitions=64×10, scan time=42min). The minimum allowed TE for each sequence was selected (TE_PRESS_=15ms, TE_sLASER_=20ms, TE_ISIS_<1ms). The overall scanning time was 2h20 per mouse. The RF power was optimized in phantoms and *in vivo* to adjust for maximal signal Intensity. Optimal TR was assessed by measuring the T1 of PCr in phantoms using 3D-ISIS with inversion recovery and by maximizing the relative SNR (PCr/yATP) in one mouse *in vivo*. Focus was put on obtaining an optimal TR that benefits the signal resonances with highest intensity (ATP, PCr and Pi).

Spectra were processed (phase and B0-drift corrections, and averaging) in jMRUI and analysed with AMARES^35^, using Lorentzian line-shape modelling and constrained frequency, linewidth and amplitude for each component (PCr, γATP, αATP, βATP, Pi_in, Pi_ex, PE, PCho, GPC, GPE, NADtot, NAD+) with additional FID weighting in the first 20 points. [Mg2+] and pH were determined as defined by AMARES, i.e. Pi_in-PCr and βATP -PCr chemical shift differences. Cramer-Rao Lower Bounds (CRLB%) were calculated by dividing the fit error (SD) obtained from AMARES quantification over the signal amplitude^36^. SNR of each scan was measured with MATLAB using the raw signal of PCr (peak height) and noise between 10-20ppm.

### Saturation transfer ^31^P-MRS

For saturation transfer ^31^P-MRS, a 210uL voxel (7×5144 cl:726mm^3^) was placed in the center of the brain. To limit the effect of contaminating signal from adjacent tissues, such as muscles, Outer Volume Suppression module was included (5mm thick slices with no gap at each voxel side). and Shimming (MAPSHIM) was performed in the voxel to reach a water FWHM of 24±2Hz, followed by a ^31^P-MRS acquisition using 3D-ISIS (Npoints=1024, AcquisitionBW=40ppm, TR=8s). Progressive saturation transfer experiment was performed using the BISTRO (B1-insensitive train to obliterate signal) method^37^, using a train of eight frequency selective 40ms HS2 pulses (BW=100Hz) at yATP offset, i.e. −2.50 ppm^38^ with variable RF field amplitude (with following scaling factors: 0.02, 0.04, 0.07, 0.14, 0.27, 0.49, 0.82, 1). Saturation time was acquired in random order at 5 different values (336, 672, 1’009, 2’018 and 4’710 ms) and a control spectrum was acquired with a mirrored saturation at +2.50ppm. To provide a narrow and consistent saturation of *γ*-ATP resonance, the effective saturation bandwidth of the BISTRO protocol was compared to similar protocols using constant flip-angles using HS2 pulses or sinc3 pulses in phantoms. Each saturation transfer experiments lasted ∼2h per subject. After spectra post-processing in jMRUI, the PCr signals were fitted using a single Lorentzian line-shape in AMARES. A mono-exponential function was fitted (MATLAB) to the relative PCr signal (M_PCr_) decay as a function of saturation time (t_sat_) using following equation:

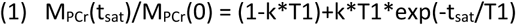

To determine both k, the pseudo first-order forward reaction PCr→ATP rate constant (k_PCr→ATP_), and T1, the apparent T1 of PCr^22^. The pH was determined from the chemical shift difference between Pi and PCr on the averaged spectrum for each mouse.

### Statistics

Statistics were all performed with GraphPad Prism 9 (GraphPad Software, San Diego, CA, USA). All values are given as mean±s.d. unless stated otherwise. P-values of *P*<0.05 were considered statistically significant.

Sequence SNRs were compared using a repeated-measure 2-way ANOVA with a Bonferroni post-hoc test. For all three sequences, the between-session coefficient of variation (CV, i.e. SD/Mean) was assessed for increasing scanning time, i.e. for increasing number of spectra averages, using a Bland-Altmann standard deviation (SD_BA_)^39^.

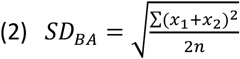

Where, x_1_ and x_2_ represent the two repeated measurements of the two sessions for the n subjects.

For 3D-ISIS, the metabolite’s CVs were computed for between-group, within-session and between-session with increasing scanning time. Between-group SD was calculated from session 1, within-session SD was calculated from the average quantification for all mice (session1+2) over the time of acquisition for 3 different time resolutions (4, 8 and 13min), and finally the between-session CV was determined using a Bland-Altmann SD (SD_BA_).

For the power calculations, the following equation was used^40^:

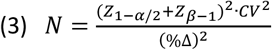

With N, the number of subjects per group, Z scores for α=0.05 and β=20% (*Z*_1–*⍺*/2_ = 1.96 and *Z*_*β*–1_ = 0.8416), CV the coefficient of variation and %Δ the precent change expected.

Fitting of the creatine CK rate constant k_f_ was done using MATLAB (R2020a) using a Levenberg-Marquart algorithm. Correlations between k_f_ and other parameters were estimated using Pearson correlation coefficients. Group comparisons between high- and low-respiration rates were done using Student’s t-tests.

## Results

### Localized ^31^P-MRS in mouse brain at 7T using 3D-ISIS leads to higher SNR and better repeatability than PRESS and sLASER

For each mouse, a set of spectra was acquired using each of the three ^31^P-MRS sequences i.e. 3D-ISIS, PRESS and sLASER to assess their performance in mouse brain at 7T. Each sequence led to a measurable PCr signal (**Fig.1a-b**), with clear visual differences at the level of metabolites with short T_2_-relaxation (e.g. *γ*-, *⍺*- and *β*-ATP). The SNR of the PCr peak was measured in each spectrum (**Fig.1c**), indicating comparable values between 3D-ISIS and sLASER (n.s.) but a better performance of 3D-ISIS over PRESS (p<0.0001) in both scan sessions. Finally, to test the repeatability of each sequence, the coefficient of variation (CV) of the SNR was calculated when increasing the number of included averages of each scan (**Fig.1d**). This analysis indicated that 3D-ISIS has the lowest CV of SNR_PCr,_ which stabilized at 15-16% for acquisitions >20 min of scan. Interestingly, while sLASER performed better for short acquisitions (<20min) with a CV under 20%, its repeatability deteriorated for longer scans, suggesting potential susceptibility to motion or other artifacts during prolonged scans.

**Figure 1:**
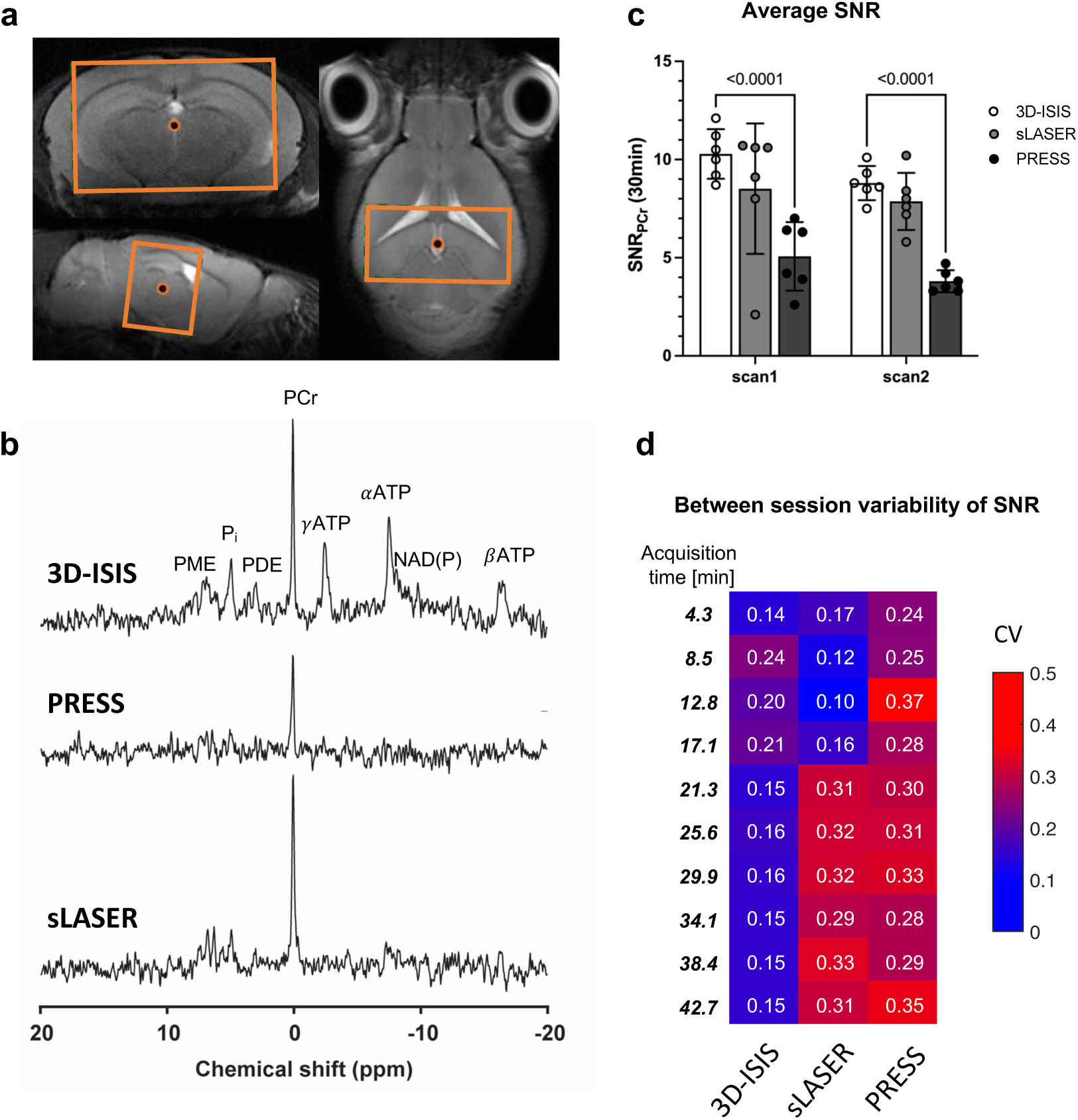
Sequence comparison for localized ^31^P-MRS in mouse brain at 7T. **(a)** Anatomical T2-weighed MRI with voxel placement (orange) in the mouse brain (8×5175 cl:1064 mm^3^). **(b)** Example of ^31^P-MRS spectra acquired using 3D-ISIS, PRESS and sLASER in the same mouse. Spectra are shown with 10Hz apodization. PCr: phosphocreatine; (*γ*, *⍺* and *β*)ATP: gamma, alpha and beta-adenosine triphosphate; P_i_ : inorganic phosphate; PME: phosphomonoesthers; PDE: phosphodieters; NAD(P): Nicotinamide andenine dinucleotide (Phosphate); **(c)** Average SNR (PCr peak) for each scan session. Sequence effect (F(2,15)=26.14, P<0.0005), p-value for Bonferroni post-hoc test is shown. **(d)** Coefficients of variation (CV) of the SNR (PCr) of each sequence at increasing number of averages or acquisition time.

### Acquisition with 3D-ISIS leads to reliable and repeatable quantification of most ^31^P-metabolites at 7T

We then looked at how well ^31^P-containing metabolites could be quantified using the 3D-ISIS sequence (**Fig2.a-b**). The quantification error of high-signal resonances, such as phosphocreatine (PCr), inorganic phosphate (Pi), and adenosine triphosphate resonances (*γ*-, *⍺*- and *β*-ATP), dropped and stabilized after the first 20min of scan, leading to relatively low Cramer-Rao Lower Bounds (CRLB): CRLB(PCr) = 7-9%, CRLB(*γ*-ATP) = 21-30%, CRLB(*⍺*-ATP) = 25-33%, CRLB(*β*-ATP) = 34-42%, and CRLB(Pi) = 45-54%. Within 45 min of acquisition, the CRLB of phosphoethanolamine (PE), glycerophopshorylcholine (GPC) and total nicotinamide andenine dinucleotide (NAD_tot_) also dropped and stabilized: CRLB(PE) = 54-65%, CRLB(GPC) = 66-77% and CRLB(NAD_tot_) = 74-94%. Finally, the CRLBs of glycerophosphorylethanolamine (GPE), NAD^+^, and phosphorylcholine (PCho) were all above 100% within 45min of acquisition in the 160uL voxel indicating poor quantification performance. Importantly, the data quality did not allow to distinguish the intracellular (Pi_intra_) component (at 4.9 ppm) from the extracellular (Pi_extra_) component (at 5.3 ppm) of inorganic phosphate when fitting them separately.

**Figure 2:**
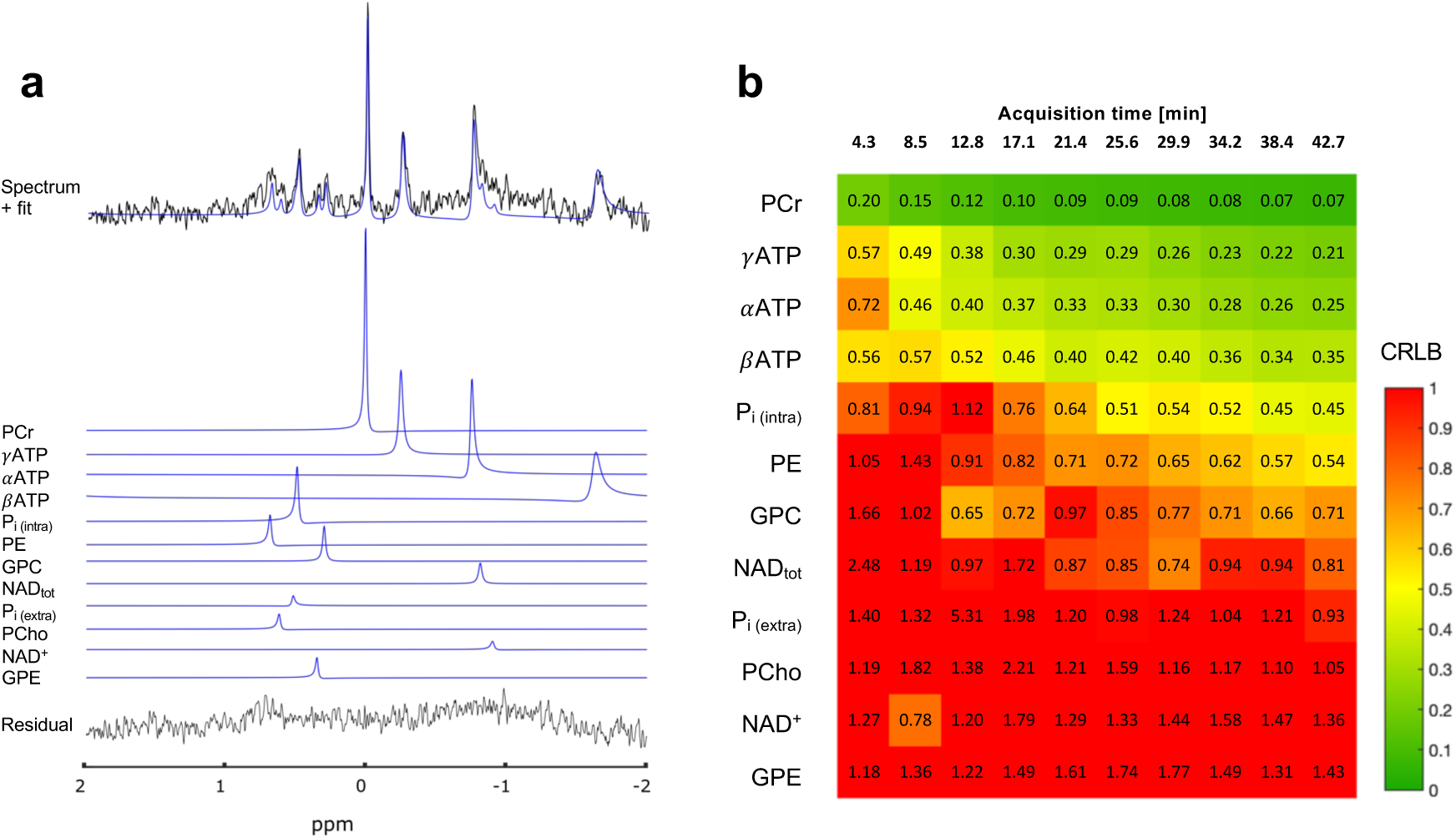
Quantification and measurement errors of ^31^P-MRS using 3D-ISIS. **(a)** Typical 3D-ISIS spectrum (black) from 160uL voxel and fit (blue) from AMARES (jMRUI). **(b)** Average quantification error (CRLB) resulting from AMARES fit for each individual component for increasing acquisition time (number of spectra averages). PCr: phosphocreatine; (*γ*, *⍺* and *β*)ATP: gamma, alpha and beta-adenosine triphosphate; P_i(intra)_: intracellular inorganic phosphate; PE: phosphoethanolamine; GPC: glycerophosphorylcholine; NAD_tot_: Nicotinamide andenine dinucleotide (1/2NAD(P)^+^+NAD(P)H); P_i(extra)_ extracellular inorganic phosphate; PCho: phosphorylcholine; NAD^+^: Nicotinamide andenine dinucleotide; GPE: glycerophosphorylethanolamine.

High CRLB values are not necessarily suggesting poor quantification, as low concentration metabolites will inevitably have higher CLRBs than high concentration ones^41^. To take this effect into account, we also looked at the CV of the measured concentration (relative to PCr) for each metabolite with increasing number of averages (**Fig.3a-c**) as a measure of quantification reliability. Results for ‘between group’ comparison indicated that Pi/PCr and (*γ*-, *⍺*- and *β*-)ATP/PCr reached a CV under 50% within 20min of acquisition. Among all other metabolites, only NAD_tot_/PCr and PE/PCr showed a decrease in CV reaching <50% within 40min of acquisition (**Fig.3a**). Between-session variation was comparable for ATP/PCr but worse for Pi/PCr, despite still being under CV=50% within 40min of acquisition (**Fig.3b**). Among small concentration metabolites, only NAD_tot_ showed a drop of CV with acquisition time, while all others were above or close to CV=50%. When looking at within-session comparison, all CVs were under 40%, with half of the metabolites falling under 10% for an 8 min spectral average (**Fig.3c**). Power calculations were then performed using these CV, on the highest signals, i.e. *γ*-ATP/PCr and Pi/PCr, to estimate the potential sample size of animals required for three experimental designs (i.e. between-group, between-sessions and within-session) (**Fig.3d-f**). Power calculations for a within-session comparison suggest that changes of ATP/PCr ratio above 25% could be easily detected for a 13 min timepoint resolution using around 10 animals. This confirms that reliable measurement of changes in PCr signal during a saturation-transfer experiment with a spectral average acquisition of 13 min for each saturation time can be achieved.

**Figure 3:**
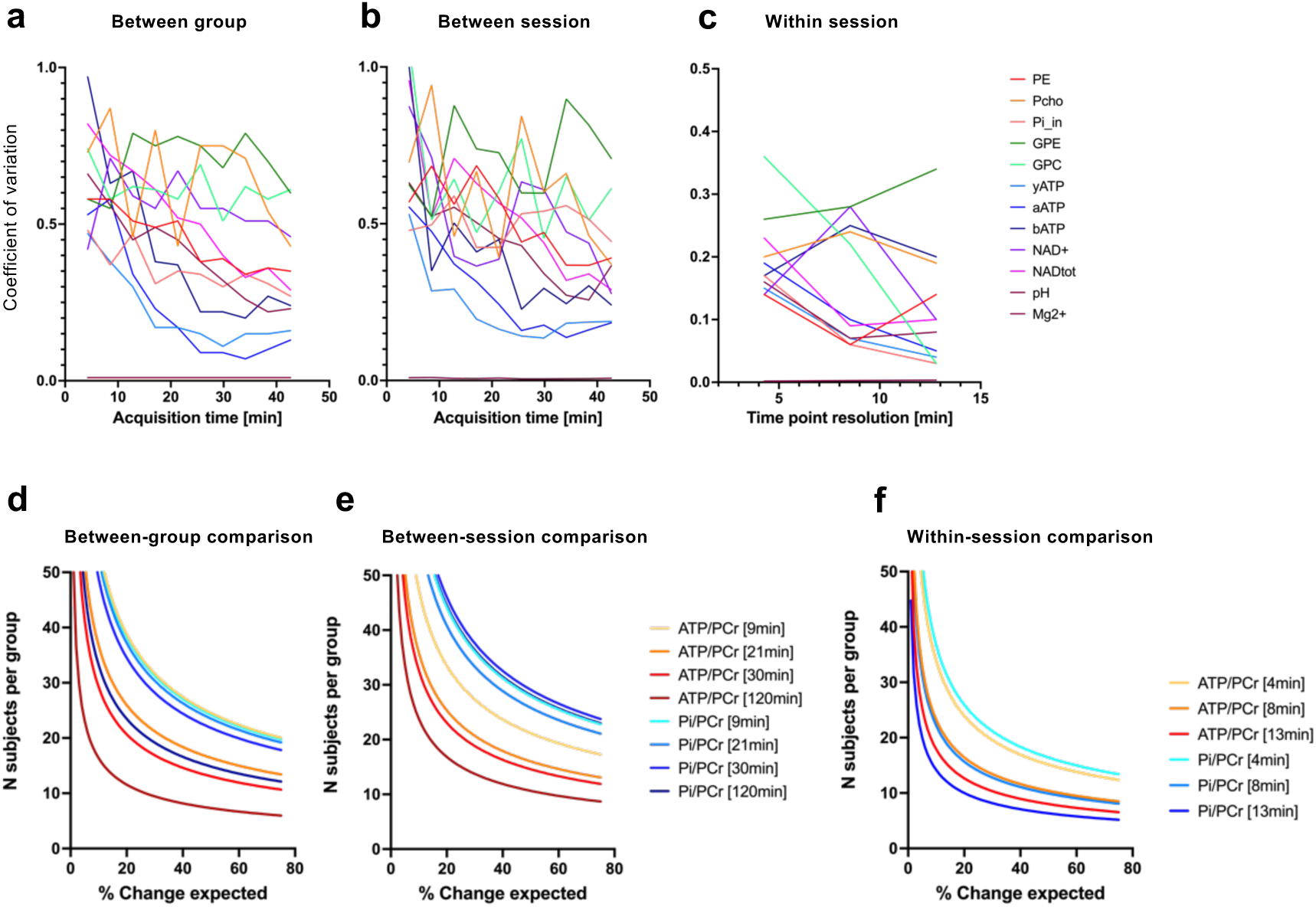
Test-retest analysis of individual ^31^P-MRS metabolites for repeatability assessment and power calculations. **(a-c)** Coefficient of variation (CV) for each metabolite as a function of acquisition time (number of spectra averages), when considering a two-group comparison (a), two-session repetition (b) and within-session fluctuation. **(d-f)** Associated statistical power calculations for ATP/PCr and Pi/PCr ratios at different acquisition times. The CV at 120min was estimated from extrapolating the CV decay in Fig.3a-b.

### Saturation transfer (ST) ^31^P-MRS is feasible for measuring the forward creatine kinase rate constant in mouse brain at 7T

We then explored the feasibility of measuring CK activity in mouse brain using a progressive-saturation transfer experiment. This was performed in a bigger voxel (210uL, whole brain) to maximize PCr and ATP signals. BISTRO pulse train led to a clean saturation with steady saturation bandwidth as measured in phantom, compared to the two other protocols, even at long saturation times (**Fig.4a**). This resulted in an efficient saturation of *γ*-ATP resonance and drop of PCr with minimal or no bleed over in the *in vivo* experiments (**Fig.4b**).

**Figure 4:**
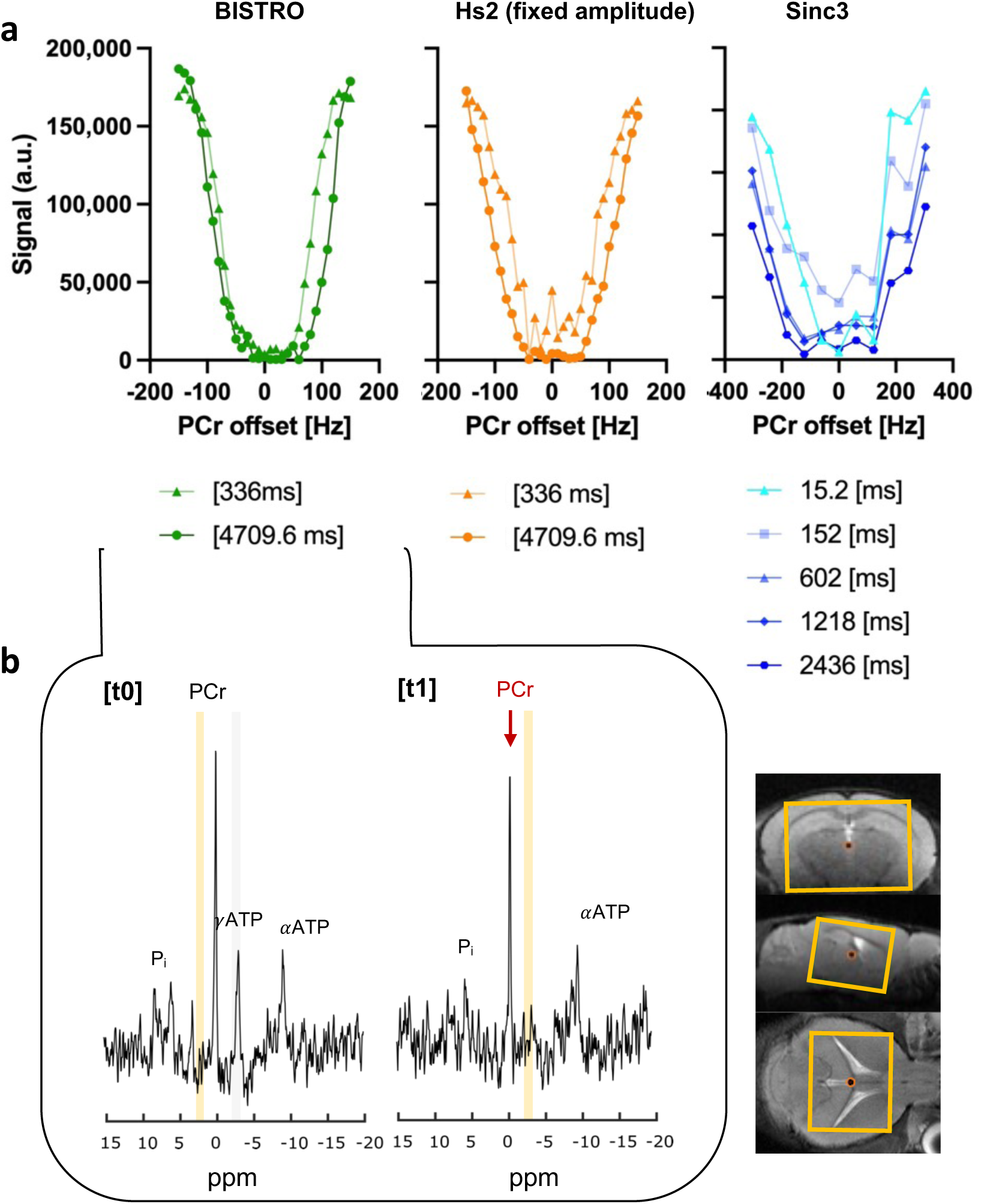
BISTRO optimization for ^31^P-MRS saturation-transfer in mouse brain at 7T. **(a)** Saturation pulse bandwidth comparison in phantom using BISTRO (green), HS2 pulse with fixed amplitude (orange) or Sinc3 pulse with fixed amplitude (blue) at various saturation times. **(b)** Typical ^31^P-MRS spectrum before [Sat_offset_(t_0_)=2.50ppm] and after [Sat_offset_(t_1_)=-2.50ppm] saturation of *γ*ATP in mouse whole brain voxel (orange; 7×5202 cl:4696mm^3^). Spectra shown with 10Hz Lorentzian apodization.

### ST ^31^P-MRS detects physiologically-relevant changes in brain creatine kinase function in mouse brain

Finally, to test the biological relevance and applicability of saturation-transfer ^31^P-MRS acquisitions in the mouse brain, we investigated the relationship between the CK forward rate constant (k_f_) with various physiological, biological and acquisitions-related parameters. The pseudo first-order CK forward rate constant (k_f_; **Fig.5a**) was determined for each mouse by fitting an exponential decay on the PCr saturation time course. The resulting k_f_ were then correlated with the various parameters and revealed a strong correlation with T_1_ (R=-0.90, p<0.0001). This indicates a strong dependence of these two variables, suggesting difficulty to infer them independently from the fitting approach (**Fig. 5b**). Nevertheless, k_f_ were also correlated with the respiration rate (R=-0.56, p=0.01) and the pH (R=-0.49, p=0.047) indicating a physiologically relevant relationship. Alternatively, no significant correlations were found for body weight, age, isoflurane dose, temperature and SNR. Importantly, k_f_ did not significantly correlate with the metabolite ratios *γ*-ATP/PCr and Pi/PCr, supporting the idea that the metabolite concentrations remain in steady-state independently of the CK flux. No group differences in k_f_ were also observed between male and females (n.s.).

**Figure 5:**
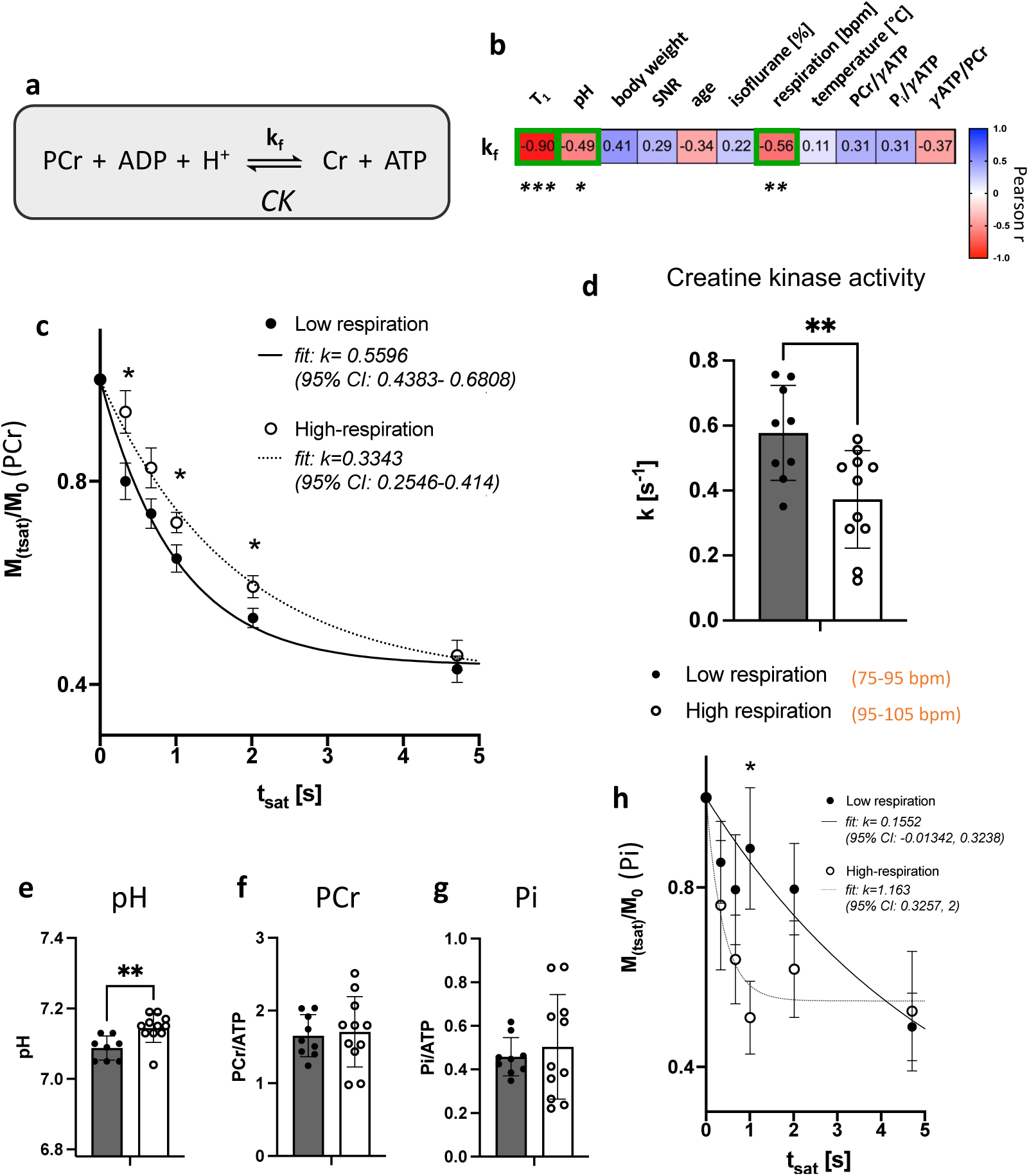
Breathing rate-dependent changes in CK activity can be measured in mouse brain using saturation-transfer experiments at 7T. **(a)** Chemical reaction catalysed by creatine kinase (CK). k_f_: CK pseudo first-order forward rate constant. **(b)** Correlations of k_f_ with physiological, biological and acquisition-related parameters (Pearson correlation coefficient) ***p<0.0001, **p=0.013, *p=0.047. **(c)** Group average of the relative PCr signal (Mt_sat_/M_0_) with increasing ATP saturation times (t_sat_) for high- vs. low-respiration rate, following progressive saturation transfer experiment. Data is shown with exponential fit for k_f_ (group average) with 95% confidence interval (CI). Student t-test for individual time points, *p<0.05. **(d-g)** Group comparisons of k_f_ **(d)**, pH **(e)**, phosphocreatine **(f)** and inorganic phosphate **(g)** for mice with high- vs. low-respiration rate. **p<0.01 **(h)** Group average of the relative Pi signal (Mt_sat_/M_0_) with increasing ATP saturation times (t_sat_).

Respiration rate, determined by the anesthetic depth, could potentially cause changes in CK activity. We thus further tested whether high-versus low-breathing rates can give rise to distinct k_f_ values. When fitting the exponential decay model to the group average PCr-saturation data, we observed a lower k_f_ for the low-respiration group (k=0.33[s^−1^], 95% CI: 0.25-0.41[s^−1^]) compared to the high-respiration group (k=0.60[s^−1^], 95% CI: 0.44-0.68[s^−1^]). When fitting the exponential decay model to the PCr-saturation data of each mouse individually, we found that animals with lower respiration rate (75-95 bpm) had significantly lower k_f_ values (0.37±0.15[s^−1^] vs. 0.58±0.15[s^−1^], 36% decrease, p=0.007) compared to those with high respiration rate (95-105 bpm). This group comparison also revealed smaller pH values in the low respiration group (p=0.005), while no difference was observed in PCr/ATP and P_i_/ATP. CK forward rate constant is sensitive to the local concentration of protons (H^+^), thus a drop in pH is expected to lead to an increase in the measured k_f_ (**Fig.5a**). We also investigated the potential effects on the ATP-synthase rate by looking at the relative decay of Pi signal following the same ATP saturation scheme. When fitting an exponential decay to the average M_sat_/M_0_ time courses of each group, we observed a lower rate of ATP synthase in mice with low respiration (k_f_=0.15; 95% CI: −0.01-0.32) compared to high respiration (k=1.16; 95% CI: 0.33-2.00).

Taken together, these results suggest that in the anaesthetized mouse brain, the creatine kinase reaction is influenced by the breathing rate, likely due to pH changes resulting from a drop in mitochondrial ATP synthesis.

## Discussion

This study is the first to investigate the feasibility and repeatability of using ^31^P-MRS to study mouse brain metabolism with a 7T scanner. We demonstrate that, under the proposed protocol, the quantification of most ^31^P-metabolites is consistent and robust, enabling a reliable assessment of CK activity in the mouse brain. Using this methodology, we identify key neuroenergetic fluctuations that occur in conjunction with physiological changes in the animal under isoflurane anesthesia *in vivo*.

Detecting of ATP and Pi resonances in ^31^P-MRS can be challenging due to their relatively short transverse relaxation times (T_2_)^42^. Although accurately measuring T_2_ can be difficult, studies have observed a trend toward reduced T_2_ values with increasing scanner field strength^43,44^, suggesting that 7T scanner might allow for longer echo-times (TE) compared to higher field scanners (>7T). To explore this, we compared three sequences that differ in their acquisition schemes, including TE. ISIS, which is based on a multi-acquisition of free induction decay, allows for the shortest delay between spin excitation and signal acquisition^34^, making it sensitive to fast T_2_-relaxing metabolites. However, its reliance on spectral subtraction for localization makes it inherently susceptible to motion artifacts^45^. On the other hand, conventional single-shot acquisition methods such as PRESS or sLASER result in longer TE (15-20ms in our case), but may offer more stable signal acquisition, potentially better suited for long scans. sLASER, in particular, provides improved spatial localization and reduces chemical shift displacement artifacts^33^. While inhomogeneous radiofrequency (B1) fields can be problematic with surface coils, the use of adiabatic pulses in sLASER achieves more uniform and efficient signal refocusing, leading to narrower spectral lines and improved resolution. Despite producing comparable PCr signal, our results indicate that sLASER still struggles to detect short T_2_ metabolites at 7T. Notably, the CV for the PCr signal between-sessions was higher with sLASER than with 3D-ISIS, particularly over longer periods, indicating reduced signal stability.

The 3D ISIS sequence provided a reliable quantification of key energy-related ^31^P-metabolites in the mouse brain at 7T. Metabolites with clearly identifiable peaks, such as ATP, PCr and Pi achieved satisfactory quantification after 20 minutes of acquisition. Our results also suggest that some low-concentration metabolites with high CRLBs, such as NAD_tot_, PE and GPC can be measured convincingly within an hour. CRLB estimate the reliability of the quantification^46^, but interpreting them as a percentage of the estimated concentration can be misleading, as higher CRLB values do not necessarily indicate poor fitting^41^. Instead of focusing solely on absolute CRLB values, we also considered whether CRLBs decreased with increased SNR when acquisition time, or the number of spectral averages, is increased. We further assessed whether increased number of averages improved the repeatability (CV) of the metabolite quantifications. However, some metabolites, such as PCho and GPE, remained unreliable with consistently high CRLBs and CVs. Additionally, we were unable to distinguish between NAD+ and NADH or between intra- and extracellular Pi pools. Increasing B_0_ field strength reduces the spin-lattice (T_1_) relaxation time of ^31^P resonances^47^ and their peak linewidths^28^ in ^31^P-MRS acquisitions. Higher filed scanners (>7T) may thus be necessary to assess these low-concentration and closely adjacent resonances.

Our power calculations suggest that typical preclinical group sizes (10-12 mice) should suffice to detect metabolite concentration changes of around 20% with standard commercially available equipment. A between-session design exhibited the highest variability between metabolic readings, likely due to physiological factors such as anesthesia-induced metabolic changes or circadian and physiological variations. However, a “within-session” design revealed minimal variability, indicating that this method effectively detects changes occurring during the scan. This protocol is therefore well-suited for saturation-transfer experiments.

Using a BISTRO saturation sequence, we were able to measure CK forward rate constant (k_PCr→ATP_) in the mouse brain. BISTRO, which uses a gradual increase in the saturation RF pulse power, creates a sharp saturation profile that reduces bleedover on adjacent resonances, especially critical at long saturation times^48^. Fitting an exponential function to the relative PCr decay following saturation of y-ATP allowed us to infer a single k_f_ value for each mouse. However, assessing ATPase activity and fitting the model to the relative Pi signal was only possible by averaging across subjects, making group comparisons challenging. The resulting k_f_ values were all in the physiological range, (k_f_=0.46±0.18 s^−1^) and distinguished mice based on their respiration regimes. A previous study reported k_f_ values in the brain of wild-type mice at 36 weeks (k_f_=0.45±0.08 s^−1^) and 72 weeks (k_f_=0.49±0.04 s^−1^)^22^ of age. The authors found no significant group difference in kf between wild-type and 5xFAD mouse model of Alzheimer’s disease, despite an overall inverted trend in k_f_ values (0.49±0.04 s^−1^ at 36 weeks and 0.43±0.06 s^−1^ at 72 weeks). While the ages and average kf values of our cohort align with that study, the higher variability observed in our data may be due to a broader range of animal respiration rates. However, the previous study did not report individual respiration regimes, making this difficult to confirm. Finally, we observed high correlation between the two fitted parameters, T1 and k_f_, suggesting that uncertainty in their estimation may contribute to the variability observed.

We found that relatively small changes in respiration rate (10-20%) in anesthetised mice can lead to significant changes in brain energy metabolism, detectable by ^31^P-MRS at 7T. The CK-catalysed reaction (Fig.5a) helps maintain steady intracellular ATP levels when oxygen levels drop or mitochondrial ATP production decreases. The build-up of [H^+^], seen as a drop in pH due to increased glycolysis, pushes the reaction towards ADP phosphorylation via PCr through CK activity. This mechanism maintains ATP levels constant when oxidative phosphorylation is compromised. Reduced intracellular O_2_ levels or build-up of HCO_3_^−^ may drive this metabolic switch resulting from lower animal breathing rate.

Various physiological changes, such as hypoxia, hypoglycemia or temperature variations, have been reported to influence CK rate^49,50^. Brain maturation has also been associated with increased in CK activity, particularly the mitochondrial isoenzyme^51,52^. Regional differences in CK activity have been observed, with higher rates in gray matter compared to white matter^53^. CK fluxes are also affected by age, psychiatric disorders, and neurodegenerativee diseases^54,55^. Studies using ^31^P-MRS on rat brains have highlighted the influence of anesthetic regime and dosage on the overall brain energy production. For example, Bresnen & Duong reported a decrease in CK rate constant with higher isoflurane doses (1.2% vs. 2%)^56^. Sauter & Rudin found that kf correlates with brain EEG activity, with values ranging from 0.10 to 0.70 [s^−1^] when modulating brain activity with bicuculline or thiopental under halothane anesthesia^57^. Du et al. observed that deeper anesthesia in rats leads to lower ATPase and CK activities due to reduced CMR_glc_ and CMRO_2_^58^. Importantly, they did not observe a drop in brain tissue pH due to the anesthetic regime. While these studies did not examine the effects of respiration rate, they also focused on the rat, which require less frequent lung ventilation than mice. Consequently, ^31^P-MRS measurements in rats may be less sensitive to fluctuations in breathing rate than in mice. While a switch from oxidative respiration to glycolysis in mouse brains may not significantly change ATP or other metabolites that maintain homeostasis, maintaining a constant breathing rate in anesthetized animals is crucial to minimizing variability in brain energetic organization. Our results underscore the challenges of extrapolating findings from rat studies to mouse models due to significant physiological and metabolic differences between the two species.

### Future perspectives and limitations

Mouse models are essential tools in biomedical research, contributing significantly to our understanding of brain function and dysfunction in human diseases. This study provides lays the groundwork for future developments aimed at improving the applicability and interpretation of ^31^P-MRS in small rodents. Further sequence optimizations should be explored, such as implementing adiabatic excitation pulses or use of faster saturation transfer protocols (e.g. FAST^59^). Future studies characterizing the different parameters that influence CK activity in the mouse brain, such as using different anesthetics or controlling blood gases and glycemia, will help us better understand the neuroenergetic processes underlying mouse brain function.

### Conclusion

This study confirms that 7T MRI scanners can achieve reliable measurements using the proposed ^31^P-MRS protocol for both metabolite peaks and creatine kinase kinetics quantifications in the mouse brain. The observed relationship between respiration and brain k_f_ highlights the importance of careful monitoring the animal’s physiology under isoflurane anesthesia.

## Acknowledgements

This study was supported financially by the Swiss National Science Foundation (P500PM_203208 to AC). The Wellcome Centre for Integrative Neuroimaging is supported by core funding from the Wellcome Trust (203139/Z/16/Z and 203139/A/16/Z). We are also grateful to William Clarke and Ladislav Valkovic for their expertise in ^31^P-MRS and their fruitful discussions and advice. We also thank Chris Randell and his team from Pulseteq Ltd. for providing the necessary support and resources regarding the mouse brain ^31^P-coil.

## Authors’ contributions

AC designed the study. MT, AC and SS optimized the protocol. AC acquired and analyzed the data. AC and MT interpreted the data. AC drafted the manuscript. MT, AC, SS and JL assisted in revising the manuscript and approved the final version.

## Ethics Declaration

The authors declare no conflicts of interest with respect to the research, authorship, and/or publication of this article. This research was funded in part, by the Wellcome Trust (203139/Z/16/Z and 203139/A/16/Z). For the purpose of open access, the author has applied a CC BY public copyright licence to any Author Accepted Manuscript version arising from this submission.

